# Homoeologue expression insights into the basis of growth heterosis at the intersection of ploidy and hybridity in Cyprinidae (*Carassius auratus* red var. × *Cyprinus carpio*)

**DOI:** 10.1101/031096

**Authors:** Li Ren, Wuhui Li, Chenchen Tang, Jun Xiao, Xiaojun Tan, Min Tao, Chun Zhang, Shaojun Liu

**Author notes:** Corresponding author: Professor Shaojun Liu, Key Laboratory of Protein Chemistry & Developmental Biology of State Education Ministry of China, College of Life Sciences, Hunan Normal University, Changsha 410081, China. These authors contributed equally to this work (Li Ren, Wuhui Li).

## Abstract

Hybridization and polyploidization are considered important driving forces that form new epigenetic regulations. To study the changing patterns of expression accompanying hybridization and polyploidization, we used RNA-seq and qPCR to investigate global expression and homoeologue expression in diploid and allotetraploid hybrids of *Carassius auratus* red var. (♀) (R) and *Cyprinus carpio* ♂) (C). By comparing the relative expression levels between the hybrids and their parents, we defined the expression level dominance (ELD) and homoeologue expression bias (HEB) in liver tissue. The results showed that polyploidization contributed to the conversion of homoeologue ELD. In addition, hybridization had more effect on the change in HEB than polyploidization, while polyploidization has been considered to have more effect on the change of global gene expression than hybridization. Meanwhile, similar expression patterns were found in growth-related genes. The results suggested that hybridization and polyploidization result in differential degrees of maternal HEB in the three tissues tested. The results of this study will increase our understanding of the underlying regulation mechanism of rapid growth in diploid hybrids and allotetraploids. The differential degrees of global expression and homoeologue expression contribute to growth heterosis in newly formed hybrids and allotetraploids, ensuring the on-going success of allopolyploid speciation.

---

Polyploidization and hybridization are fundamental processes in evolution that result in the emergence of novel genotypes from the merger of two or more different genomes (Schultz 1969; Dawley 1987; Dufresne and Hebert 1994; Salmon *et al*. 2005). Many studies have focused on global expression between the parents and hybrid offspring to determine the mechanism of expression regulation in allopolyploids (Comai 2005). This phenomenon has been described as the evolution of gene expression, which is considered useful for adaptation and speciation (Wolf *et al*. 2010). Meanwhile, two sets of homoeologous genes and duplicated pairs may lead to changes in the expressions of some genes related to phenotypic differences in allopolyploids (Zhong *et al*. 2012; Zhou *et al*. 2014). Thus, a study of homoeologue expression would provide a useful platform to investigate genomic divergence in hybrids and polyploids.

*Carassius auratus* red var. (R) and *Cyprinus carpio* (C) are the most predominant and widespread form of cyprinid fish, and contain 100 chromosomes. After selective breeding, diploid hybrid offspring (2n = 100) were produced with 50 chromosomes from R and 50 from C (Liu 2010). Fertile tetraploid hybrids (4n = 200) were obtained on a large scale by crossing F_2_ diploid hybrids (Liu *et al*. 2001). Fluorescence *in situ* hybridization (FISH) results showed that allotetraploid fish could be identified contained two sets of R and C genomes, respectively (unpublished data). The two hybrid populations that originated from R and C provide us with a platform to study the regulation of homoeologue expression by polyploidization and hybridization.

Hybrid fish are widely distributed worldwide as a result of artificial or natural interspecies hybridization. Upon crossing the interspecies barrier, the newly formed progeny display heterosis, such as fast growth. Recent studies have focused on expression level dominance (ELD) and homoeologue expression bias (HEB) to analyse gene regulation patterns and their underlying mechanisms (Rapp *et al*. 2009; Yoo *et al*. 2013; Zhou *et al*. 2015). Studies have shown that allelic interactions and gene redundancy result in heterosis in allopolyploids relative to non-coding RNA, DNA, methylation and transcriptome changes (Michalak 2009; Ng *et al*. 2012). Previous studies in teleosts hybrids were largely based on global expression (Liu *et al*. 2012; Zhong *et al*. 2012); therefore, determining homoeologue expression is a promising way to study the regulation of the underlying expression mechanisms. In particular, analysis of the regulation of sets of growth-related genes is crucial to decipher the genomic basis of growth heterosis (Zhong *et al*. 2012).

An increasing number of studies of homoeologue expression have used RNA-seq to investigate gene expression patterns between hybrids and their parents. RNA-seq is regarded as an efficient method to examine overlapping hybridization among homoeologues (Udall *et al*. 2006; Rapp *et al*. 2009; Yoo *et al*. 2013). Meanwhile, in non-model organisms, the identification of homoeologue-specific single nucleotide polymorphisms (SNPs) in the two different genomes is also useful (Pala *et al*. 2008). Homoeologue expression is then estimated by relative expression using real-time quantitative PCR (qPCR)(Pala *et al*. 2008). In this study, we combined RNA-seq and qPCR to investigate the ELD and HEB relative to hybridization (genome merger) and polyploidization (genome doubling).

To investigate changes in homoeologue expression levels related to heterosis, particularly the underlying growth regulation mechanism, we used diploid and allotetraploid hybrids of *C. auratus* red var. (♀) and *C. carpio* (♂) in our study. By comparing with the relative expression levels between the hybrids and their parents, we defined the ELD and HEB in liver tissue by RNA-seq. Meanwhile, R/C homoeologue expression silencing was identified for certain genes, revealing epigenetic changes and underlying regulation mechanisms after genome merger and genome doubling. Seven key growth-regulated genes were studied in various tissues using qPCR. The results showed that R-bias was predominant in the diploid F_1_ hybrid of *C. auratus* red var. (♀) × *C. carpio* ♂) (F_1_) and in eighteen generations of tetraploid hybrids of *C. auratus* red var. (♀) × *C. carpio* ♂) (F_18_). Our goal was to assess the magnitude and directionality of ELD and HED relative to heterosis in different ploidy level hybrids. Therefore, these data provided a novel perspective to study expression patterns of homoeologous genes under genome merger and genome doubling, and gave us an insight into the regulation mechanism that contributed to heterosis.

## Materials and Methods

### Animals

All experiments, performed from 2012–2014, were approved by the Animal Care Committee of Hunan Normal University. The Administration of Affairs Concerning Animal Experimentation guidelines stated approval from the Science and Technology Bureau of China. The methods were carried out in accordance with the approved guidelines. Experimental individuals were fed in a pool with suitable illumination, water temperature, dissolved oxygen content, and adequate forage in the Engineering Center of Polyploidy Fish Breeding of the National Education Ministry located at Hunan Normal University, China. Approval from the Department of Wildlife Administration is not required for the experiments conducted in this paper. Fish were deeply anesthetized with 100 mg/L MS-222 (Sigma-Aldrich) before dissection.

Three female individuals of diploid *C. auratus* red var. (R), diploid *C. carpio* (C), the corresponding female interspecific diploid F_1_ hybrid of *C. auratus* red var. (♀) × *C. carpio* ♂) (Liu *et al*. 2001), and female allotetraploids of *C. auratus* red var. (♀) × *C. carpio* ♂) (Liu *et al*. 2001) (2-year-old individuals) were randomly selected. Body traits (body length, height and weight) were recorded once every month (Figure S1). To measure the DNA content of the erythrocytes from the above samples, 1–2 ml of blood was drawn from the caudal vein using syringes containing 200–400 units of sodium heparin. The blood samples were subjected to nuclei extraction and 40,6-diamidino-2-phenylindole DNA-staining with cysteine DNA 1 step (Partec). The DNA contents of the erythrocytes were then detected by flow cytometry in each sample. In addition, to detect the ploidy levels of each sample, the red blood cells were cultured in nutrient solution at 25.5°C and 5% CO_2_ for 68–72 h, and then colchicine was added 3.5 h before harvest. Cells were harvested by centrifugation, followed by hypotonic treatment with 0.075M KCl at 26°C for 25–30 min, fixed in methanol–acetic acid (3:1, v/v) with three changes. Cells were dropped onto cold slides, air-dried and stained for 30 min in 4% Giemsa solution. Good-quality pictures of the metaphase spreads from 12 individuals were observed under a microscope (Figure 1) (Xiao *et al*. 2014).

**Figure 1.**
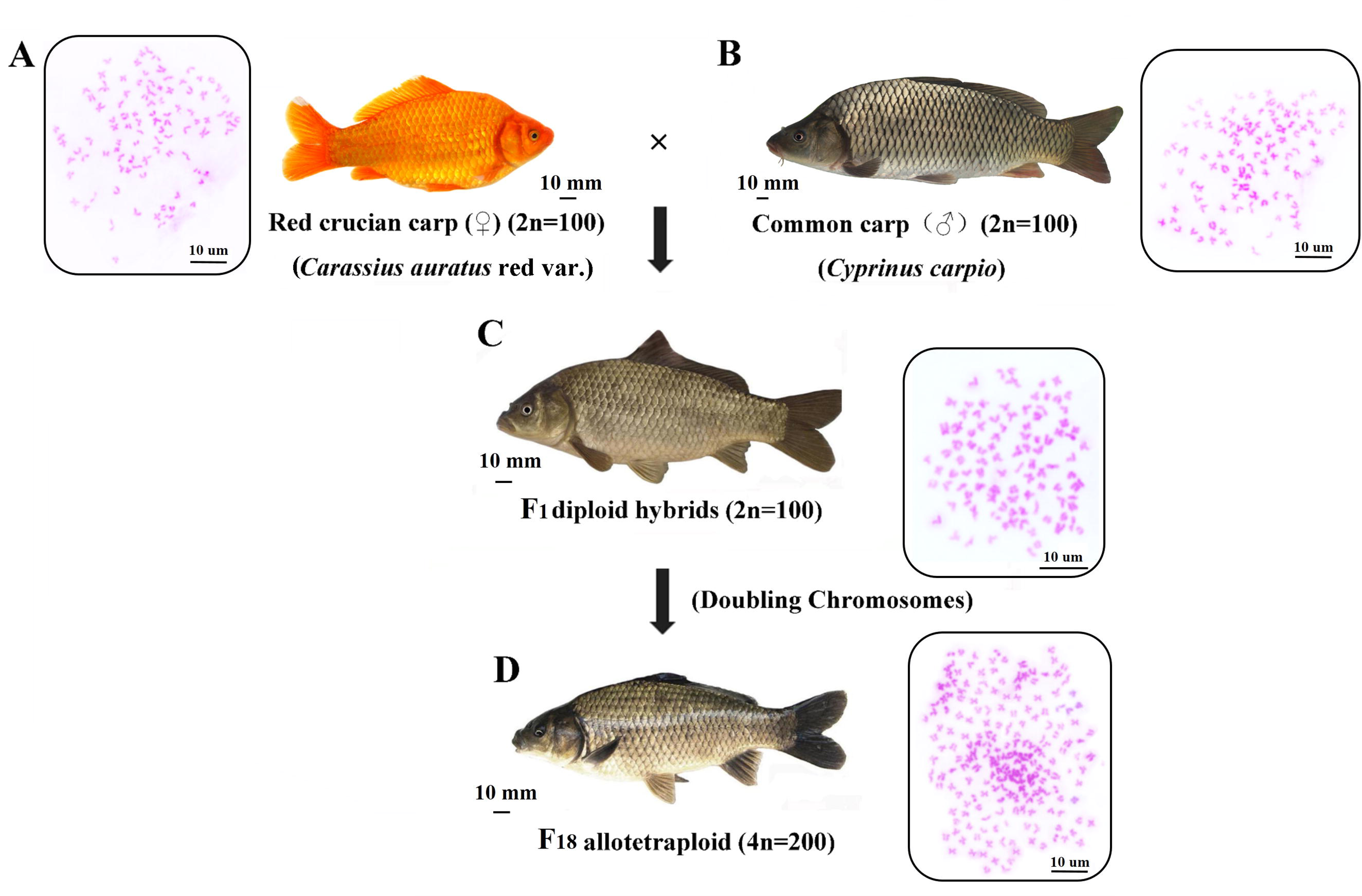
Formation of *C. auratus* red var. × *C. carpio* hybrids. A. 100 Chromosomes were observed in *C. auratus* red var. B. 100 Chromosomes were observed in *C. carpio.* C and D. After hybridization, F_1_ diploid hybrids (C) and F_18_ allotetraploid (D) were obtained. The observation of chromosomes showed that duplication of genome was occurred in the F_18_ relative to F_1_.

### Illumina sequencing

After anesthetizing the fish with 2-phenoxyethanol, liver, muscle and ovary tissues were excised and immediately placed into RNALater (AM7021, Ambion Life Technologies, Carlsbad, CA, USA) following the manufacturer’s instructions, for storage. Total RNA was extracted from three tissues after the RNALater was removed. RNA was isolated according to the standard Trizol protocol (Invitrogen) and quantified with an Agilent 2100 Bioanalyzer (Agilent, Santa Clara, CA, USA).

Twelve cDNA libraries representing each individual fish were constructed using 2 μg of mRNA. Each library was sequenced using an Illumina HiSeq™ 2000/2500. The read adaptors and low quality reads were removed from the raw reads and the clean reads from each library were examined using software FastQC (version 0.11.3). Principal component analysis (PCA) of the twelve liver transcriptomes was applied to examine the contribution of each transcript to the separation of the classes (Anders and Huber 2010; Reeb and Steibel 2013).

### Mapping and differential expression

After data quality control, fastq formatted reads from the two diploid parents and two hybrid offspring were mapped to the reference genome using TopHat2 (Trapnell *et al*. 2012; Kim *et al*. 2013). We utilized the *C. auratus* red var. genome assembly (http://rd.biocloud.org.cn/) (39,069 transcripts) and the *C. carpio* genome assembly (http://www.carpbase.org/) (52,610 transcripts) as the reference genomes because these transcripts databases were built from genome sequencing (Table S1). To identify putative orthologues between R and C, the two sets of sequences were aligned using the reciprocal BLAST (BLASTN) hit method, with an e-value cut off of 1e^−20^ (Blanc and Wolfe 2004). Two sequences were defined as orthologues if each of them was the best hit of the other and if the sequences were aligned over 300 bp. After identifying SNPs between the R and C orthologues, we mapped our reads from R and C to compare the mapping results. Reads with SNPs that differed between the R- and C-genome in the progenitors were parsed into R and C homoeologue-specific bins using custom perl scripts.

To calculate expression levels, the replicates were normalized using Cufflink (version 2.1.0) (Trapnell *et al*. 2012) and then, using the overall expression levels of both homoeologues of a gene, differential expression was assessed between the different polyploid level relative to their diploid parents, using Fisher’s exact tests (Wang *et al*. 2010). The mapping results were analysed with the DEGseq package in the R software version 2.13 (R Foundation for Statistical Computing, Vienna, Austria) (Wang *et al*. 2010). To remove the negative effect of expression noise, we restricted the analysis to genes have read counts (≥ 1) in all biological replicates. The abundance or the coverage of each transcript was determined by read counts and normalized using the number of reads per kilobase exon per million mapped reads (RPKM) (Mortazavi *et al*. 2008). The RPKM value of the reads was calculated to obtain the gene expression level. The false discovery rate (FDR) was used to determine the threshold P value in multiple tests and analyses. Meanwhile, the unigenes with FDR ≤ 0.05 and fold change > 2 were considered as differentially expression genes.

### Analyses of expression level dominance and homoeologue expression bias

We identified candidate novel expressions (new expression of a gene in liver) and silencing in the hybrids according to the standards of Yoo *et al* (Yoo *et al*. 2013). The number of novel expression and silenced genes was screened in the categories of global expression and growth-related genes (Table 2 and S3 Table). We then focused on genes that were expressed in both the diploid parents and in the hybrid offspring to analyse the ELD.

**Table 1.**
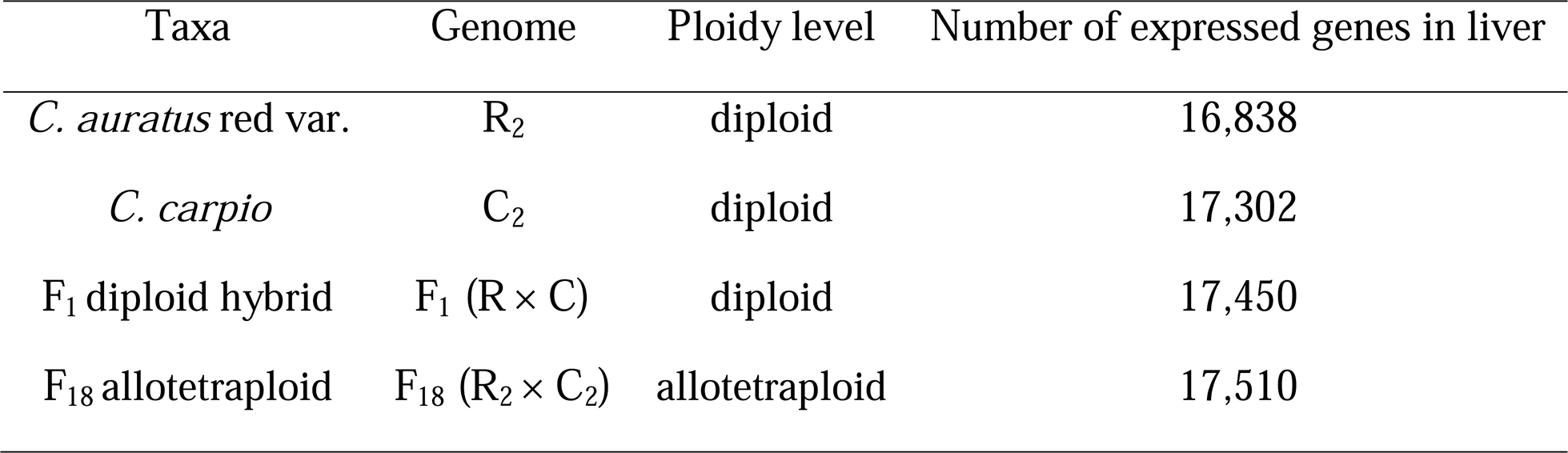
The basic information of species data used in this study

In the hybrid offspring, genes that were identified as differentially expressed in the hybrid relative to the diploid parents were binned into 12 possible differential expression categories (Figure 3), ELD, mid-parents, and up/down expression (outside the range of either parent), according to Rappet *et al*. (2009). Briefly, genes were parsed into these 12 categories (using Roman numerals; see Figure 3), depending on the relative expression levels between the hybrid and the diploid parents. In this manner, genes may display mid-parent (XI and XII), paternal C-ELD (VII and VIII), maternal R-ELD (IX and X), expression lower than both parents (I, II, and III), or expression higher than both parents (IV, V, and VI).

To describe the extent and direction of HEB in response to hybridization and evolution at the polyploid level, we analysed the differential expression across the F_1_ diploid hybrid, F_18_ allotetraploid, and the *in silico* MPVs. Values from the three biological replicates of each parent were averaged to calculate the MPV and then analysed in the same manner as described above.

### Expression of growth-related genes in RNA-seq and qPCR

Among the 3540 genes used in the study of HEB in hybrids, thirty-four growth-regulated genes were selected and analysed to help us understanding the effect from either parent based on the RNA-seq data (Table S4).

To further validate the HEB related to growth regulation in the F_1_ and F_18_, we selected seven key growth-regulated genes and subjected than to homoeologue-specific qPCR (Pala *et al*. 2008). Total RNA was extracted from the three tissues and first-strand cDNA was synthesized using AMV reverse transcriptase (Fermentas, Canada) with an oligo (dT)_12–18_ primer at 42 °C for 60 min and 70 °C for 5 min. The conserved region of the teleost orthologues’ *vasa* genes was used as a template to design universal primers (Table S5). The PCR products were cloned using appropriate primers and sequences in six parental samples and six hybrid samples. The sequences of other genes (*igf1*, *igf2, ghr, tab1, bmp4,* and *mstn*) were obtained from the assembly of liver transcriptome data.

Comparison of the sequences was done using Bioedit ver. 7.0.9, and an analysis of cDNA polymorphisms in the transcripts revealed R and C homoeologue expressed in hybrid. SNPs between the R and C homoeologues were obtained from one gonad-specific gene (*vasa*), a housekeeping gene (*β-actin*), and ubiquitously expressed gene (*igf1, igf2, ghr, tab1, bmp4,* and *mstn*). The SNP regions were used to design R/C homoeologue-primers for qPCR (Figure S6 and Table S6). The R and C homoeologue-specific primers were obtained to permit the detection of only R or C homoeologues by qPCR using the ABI Prism 7500 Sequence Detection System (Applied Biosystems, USA) (Table S7). Amplification conditions were as follows: 50 °C for 5 min, 95 °C for 10 min, and 40 cycles at 95 °C for 15 s and 60 °C for 45 s. Each test was performed three times to improve the accuracy of the results. Finally, relative quantification was performed and melting curve analysis was used to verify the generation of a single product at the end of the assay. Triplicates of each sample were used both for standard curve generation and during experimental assays. After obtaining the R and C homoeologue expression levels of the seven genes, the relative expression of each homoeologous gene was calibrated with *β-actin,* and the relative mRNA expression data were determined using the 2^−ΔΔCt^ method (Livak and Schmittgen 2001). The expression level of the reference gene *β-actin* in the hybrids was estimated using the ratio of the transcript abundance to the gene copy using PCR and qPCR of co-extracted DNA and RNA from the ovaries of diploid and allotetraploid individuals. *β-actin* expression is the same between fish of different ploidy and genome constitution, and in somatic organs and gonads (Tao *et al*. 2008; Long *et al*. 2009; Liu *et al*. 2010; Liu *et al*. 2012).

## Results

### Statistical mapping of RNA-seq data

To investigate how hybridization and polyploidization affect growth regulatory mechanism, we used the allotetraploid line of *C. auratus* red var. × *C. carpio* to study the pattern of global expression and homoeologue expression in two different ploidy level hybrids (Figure 1). The F_1_ diploid hybrid and F_18_ allotetraploid individuals were sexually mature cyprinid fish that possess hybrid traits (Liu *et al*. 2001). All short-read data have been deposited at the Short Read Archive (SRA) under accession numbers SRX668436, SRX668453, SRX671569 and SRX668467. We then annotated the exons of R and C using BLASTX alignment (e-value ≤ 1e^−6^) with protein databases (Table S1). 20,169 genes were identified in the R genome assembly and 20,365 genes in the C genome. Meanwhile, 739 million (M) clean reads (76.8%) from 12 libraries were surveyed to map to the two references sequences (Table S1 and S2). The liver transcriptome results showed that approximately 17,275 genes were expressed in four kinds of fish (Table 1). Notably, slightly more genes were expressed in the hybrids than in both of their diploid parents. This phenomenon also reflected the coexistence of R- and C-genomes in hybrid individuals.

### Differential gene expression, novel expression and silencing

To study gene expression patterns in F_1_ diploid hybrids and F_18_ allotetraploids, we performed pairwise comparisons between the diploid parents to assess pre-existing differential gene expression (Figure 2). Approximately 5104 genes (33.32%) were differentially expressed between the diploid parents (*P* < 0.05 in comparisons; Fisher’s exact test). In all comparisons, the percentage of genes showing differential expression between the F_1_ or F_18_ and their two parents was asymmetric (*P* < 0.05; Fisher’s exact tests). Meanwhile, the differentially expressed genes exhibited a bias toward the different parents. For example, global expression of the F_1_ was closer to the maternal R than to paternal C. Approximately 18.31% of genes were differentially expressed between the F_1_ and R, whereas the number of differentially expressed genes was 26.45% relative to C (*P* < 0.05 in comparisons; Fisher’s exact test). Conversely, the global expression patterns in F_18_ were closer to the paternal C than to the maternal R.

**Figure 2.**
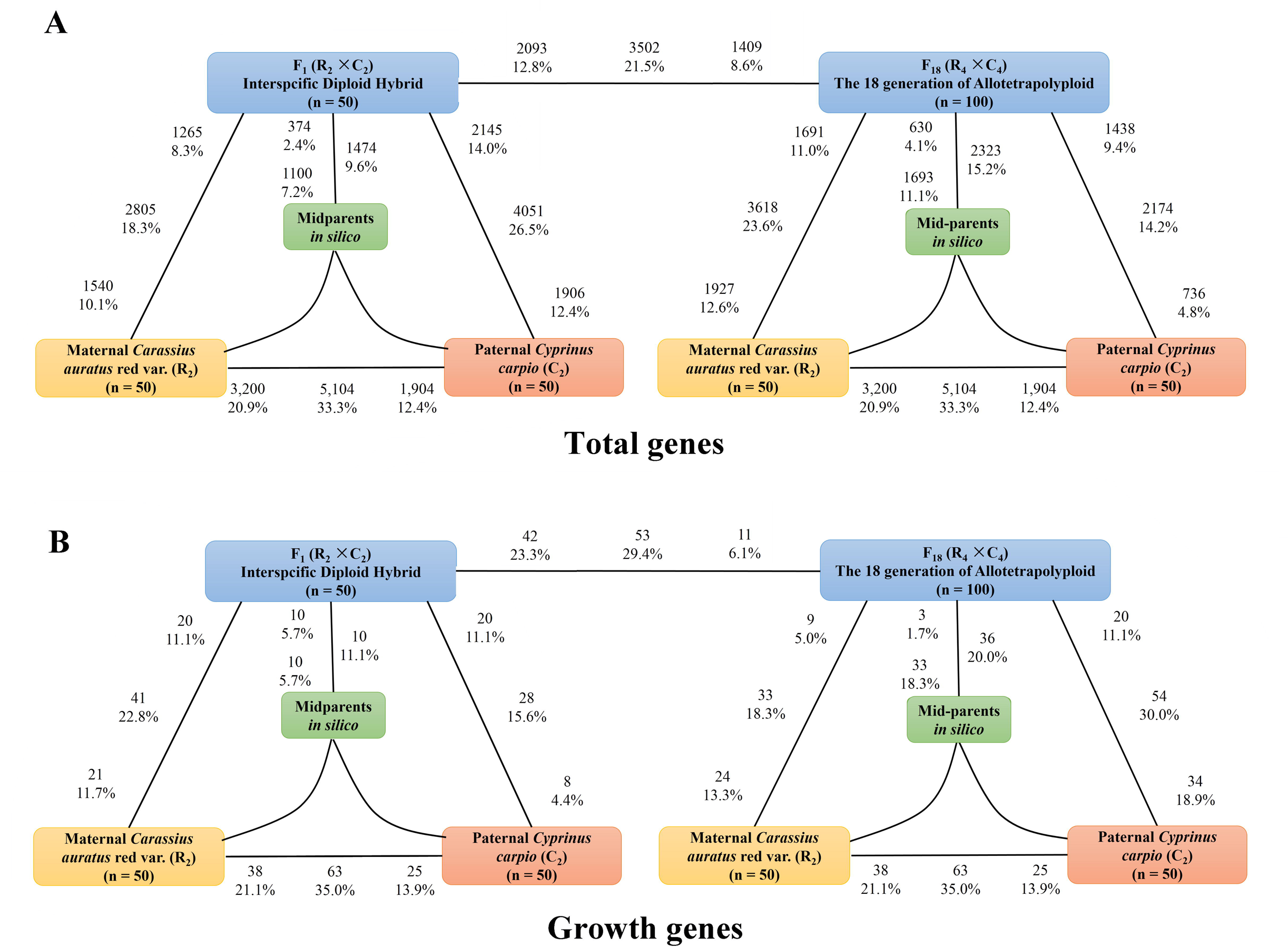
Differentially expressed genes in each contrast between hybrids offspring and their origin parents. A. Bold text exhibits the total number and fraction of genes differentially expressed in each contrast. Also shown for each contrast is the partitioning of the total number of differentially expressed genes into the direction of up regulation. For example, in (a), 5104 genes are indicated as being differentially expressed between *C. auratus* red var. and *C. carpio.* Of these, 3200 are upregulated in *C. auratus* red var., and 1904 genes are upregulated in *C. carpio.* The asymmetry between differential expression between the progeny and its diploid parents corresponds to genome-wide ELD toward one parental genome. The left figure show an interspecific diploid hybrid F_1_ generated from the diploid parents *C. auratus* red var. (R) and *C. carpio* (C). The middle of figure show that the F_18_ allotetraploid was generated from duplication of genome of F_1_. The right figure exhibits that F_18_ genome were consist of *C. auratus* red var. homoeologs and *C. carpio* homoeologs. B. Bold text exhibits the 118 growth genes number and fraction of genes differentially expressed in each contrast. Also shown for each contrast is the partitioning of the growth genes number of differentially expressed genes into the direction of upregulation.

In the expression comparison, only 13 genes (0.08%) exhibited novel expression in F_1_. However, novel expression increased with polyploidization: 44 (0.25%) genes exhibited novel expression in the F_18_ (Table 2). We then evaluated homoeologues silencing in total expressed genes in the four kinds of fish. There were 38 (0.22%) cases of R homoeologue silencing in the F_1_ and 26 (0.15%) cases in the F_18_. Nineteen (0.11%) C homoeologues were silenced in the F_1_ and 46 (0.27%) in the F_18_ (Table 2). These results suggested that polyploidization accelerates the occurrence of homoeologue silencing.

**Table 2.**
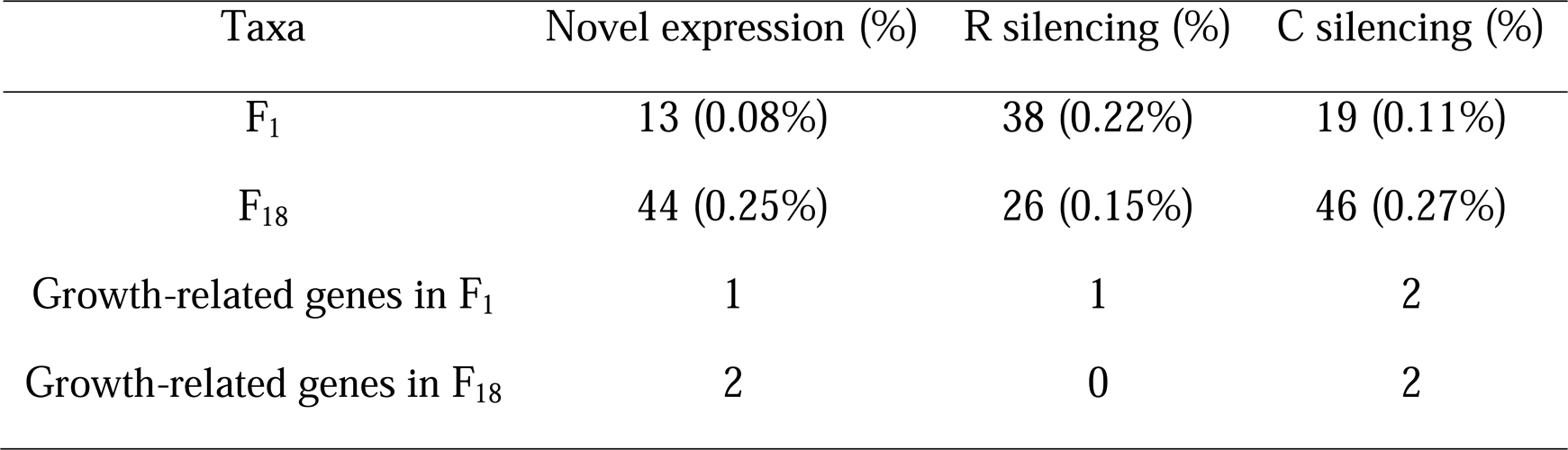
The number of genes showing the patterns of novel expression and expression silencing between the hybrids with their origin parents (at threshold of 10 reads homoeolog per million reads)

### Expression level dominance in the liver transcriptome

To study ELD in the F_1_ diploid hybrids and F_18_ allotetraploids, we performed pairwise comparisons between the hybrid offspring with the diploid parents to assess differentially expressed genes. Compared with the maternal R, 2805 (18.31%) of the F_1_ genes were identified as significantly differentially expressed, and 3618 (23.61%) such genes were identified in F_18_ (*P* < 0.05 in comparisons; Fisher’s exact test) (Figure 2). For genes pairs between the hybrid and paternal C, 4051 (26.45%) differentially expressed genes were detected in the F_1_, and 2184 (14.19%) genes in the F_18_ (*P* < 0.05 in comparisons; Fisher’s exact test) (Figure 2). To better study the ELD, we binned gene pairs from the hybrids into 12 categories including mid-parents (XI and XII), up/down expression (I, II, III, IV, V, and VI), and ELD (VII, VIII, IX and X) (see Methods). Categories VII and X represented gene pairs showing upregulated ELD in the hybrids. For example, our results showed that maternal effect played prominent role in the F_1_ (R *vs*. C = 1277 *vs*. 517), and paternal effect predominated in the F_18_ (R *vs*. C = 779 *vs*. 1061) (Figure 3). Conversely, categories VIII and IX represented the gene pairs showing downregulated ELD in the hybrids (Figure 3).

**Figure 3.**
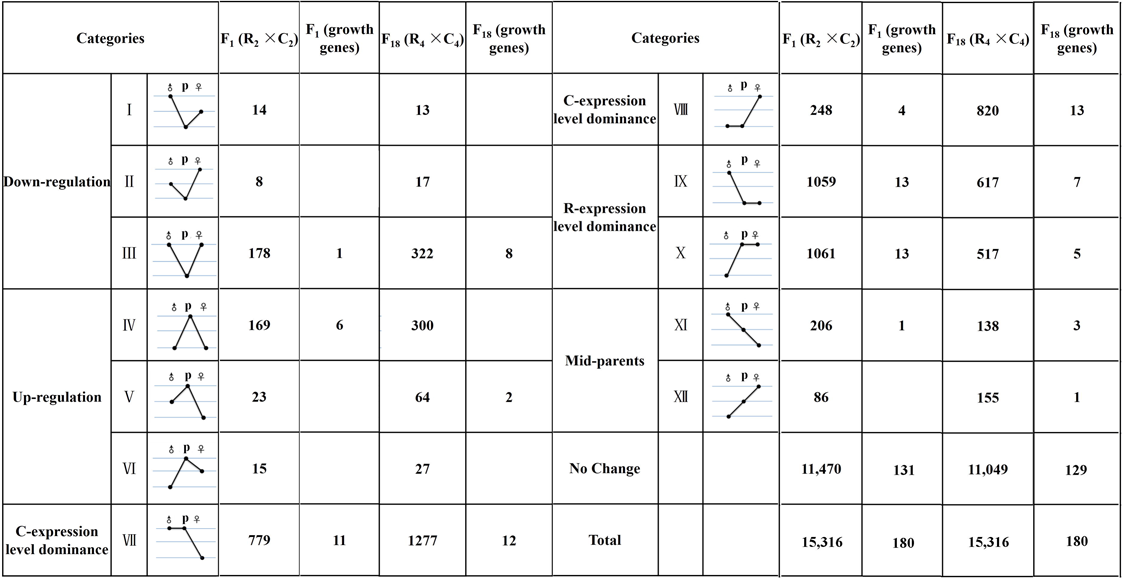
The 12 possible differential expression states in the F_1_ diploid hybrid and F_18_ allotetraploid relative to their diploid parents. Roman numerals indicate the same categorization as used in Rapp et al. (2009), with figures schematizing their respective gene expression pattern for the R-genome, diploid maternal parent (♀), the F_1_ or F_18_ (P) and the C-genome diploid paternal parent ♂).

### Homoeologue expression bias in different ploidy levels

According to the report of Rappet *et al*. (2009), the expression categorisation would not only help in the study of ELD, but also provides an insight into the HEB in the hybrids. The unbalanced gene number (VII and X *vs*. IX and X) reflected a preference toward the paternal or maternal expression in the hybrids. For example, among the 15,316 expression pairs of the F_18_, we determined that approximately 13.69% of all genes (categories VII and VIII) showed C-ELD, and 7.40% (categories IX and X) showed R-ELD, which indicated the phenomenon of C-HEB in the F_18_. Likewise, we examined the F_1_ for evidence of R-HEB, in which 2120 genes (13.84% of all genes) (categories IX and X) fell into the R-ELD category (Figure 3). Additionally, we examined the upregulated genes (IV, V, VI, X, and XII) and downregulated genes (I, II, III, IX, and XI) in the hybrids compared with the paternal C and compared the upregulated genes (IV, V, VI, VII, and XI) and downregulated genes (I, II, III, VIII, and XII) in the hybrids compared with the maternal R (Figure 3). In these comparisons, the number of significantly differentially expressed gene (up *vs*. down = 352 *vs*. 391 in F_18_, up *vs*. down = 200 *vs*. 207 in F_1_) was similar (*P* < 0.05 in comparisons; Fisher’s exact test).

To address whether the observed category of HEB really reflects the HEB in the F1 diploid hybrids and F_18_ allotetraploids, we compared 3540 genes with homoeologue-specific SNPs on a case-by-case basis between the parental diploids and their diploid hybrid and polyploids. As shown in Table 3, the patterns observed in the diploid parents were often conserved in the F_1_ and F_18_. For example, the first three rows in Table 3 show that the parental expression patterns were maintained for greater than half of all genes in this analysis: 74.8% (in F_1_) to 77.6% (in the F_18_) (*P* < 0.05 in comparisons; Fisher’s exact test). Rows 4 and 5 represent the second most common class of genes, representing 13.9–15.4% of the 3540 genes. In these cases, pre-existing expression bias in the parental homoeologues reverted to non-differential expression of the homoeologous copies in the diploid hybrids and allotetraploids (*P* < 0.05 in comparisons; Fisher’s exact test). A small numbers of genes were detected as having novel patterns that accompanied the genome merger or doubling. These cases suggested novel regulatory and/or evolutionary interactions in the hybrid offspring. We also collected genes with significant HEB in the F_1_ and F_18_ (rows 11 and 12) (Table 3 and Fig 4). In addition, to further detect the R-/C-biased in hybrids, we assessed the potential bias based on the ratio of R/C homoeologue expression levels (Table 3 rows 13 and 14). These genes helped us to understand the origin of some of the genetic traits in the hybrid offspring.

**Table 3.**
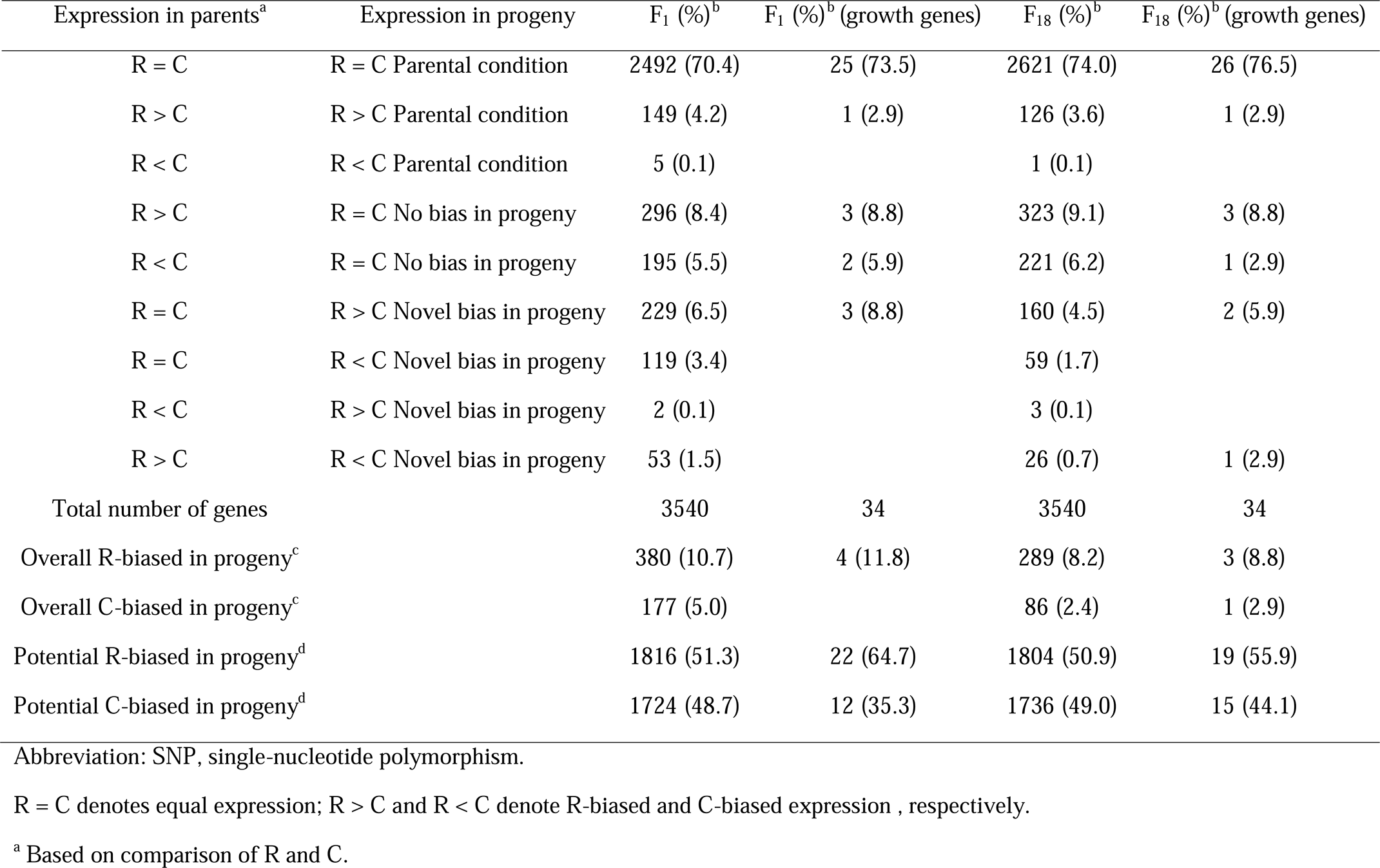
Homeeolog expression bias in the F_1_ hybrid and F_1_g allotetraploid

**Figure 4.**
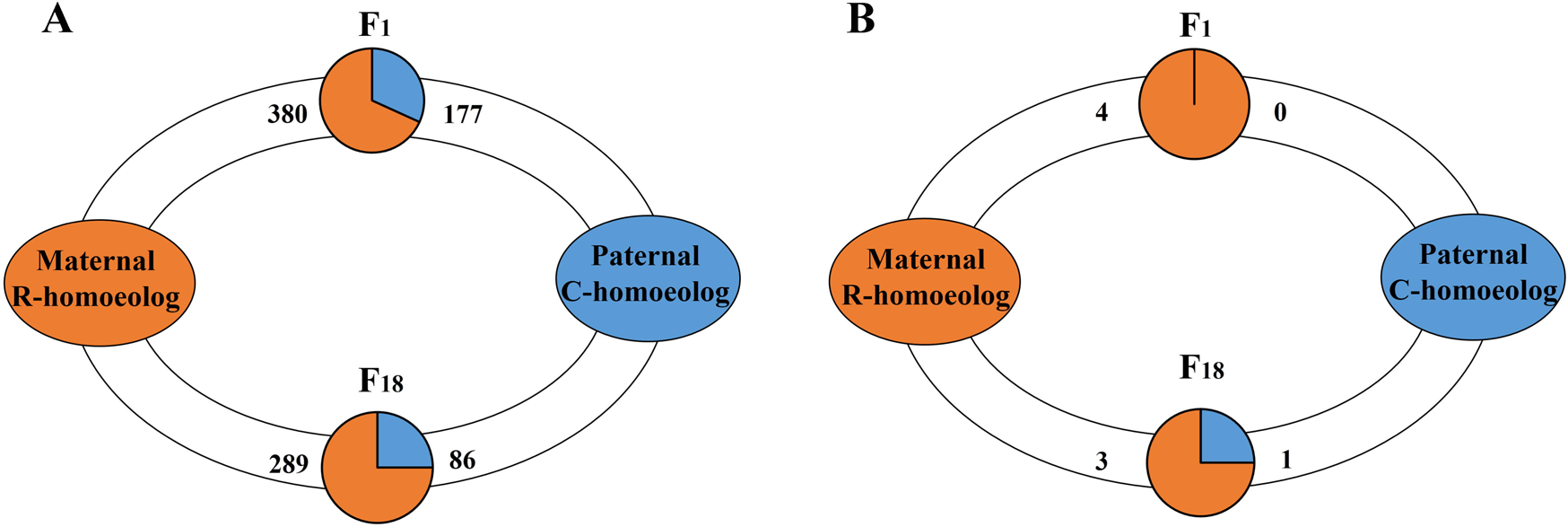
Homoeolog expression bias in total genes and growth-regulated genes of two hybrid offspring. A. The maternal HEB in total genes is estimated by the gene number of R homoeolog to C homoeolog in F_1_ and F_18_. B. The maternal HEB in growth-regualted genes is estimated by the gene number of R homoeolog to C homoeolog in both of hybrid offsprings.

For the 15,316 gene expressed in F_1_, F_18_ and their original parents, we analysed the differential expression between the hybrids with *in silico* mid-parent expression values (MPV) that replaced the expression level of both of the parents. The three categories comparison showed that only 2.8% of the genes (430 out of 15,316 genes) changed their expression patterns in response to genome merger (Table 4). As a result of genome doubling, 1893 (12.4%) genes changed their expression patterns. The results showed that genome doubling had more effect on global expression changes than the genome merger. Among the 3541 homoeologue-specific SNPs-containing genes, 75.09% (2659 genes) show no change in expression level compared with the R/C patents. However, among those that did change, the genome merger resulted in more genes with changed expression levels (13.9%) compared with genome doubling (7.4%) (*P* < 0.05 in comparisons; Fisher’s exact test, Table 4).

**Table 4.**
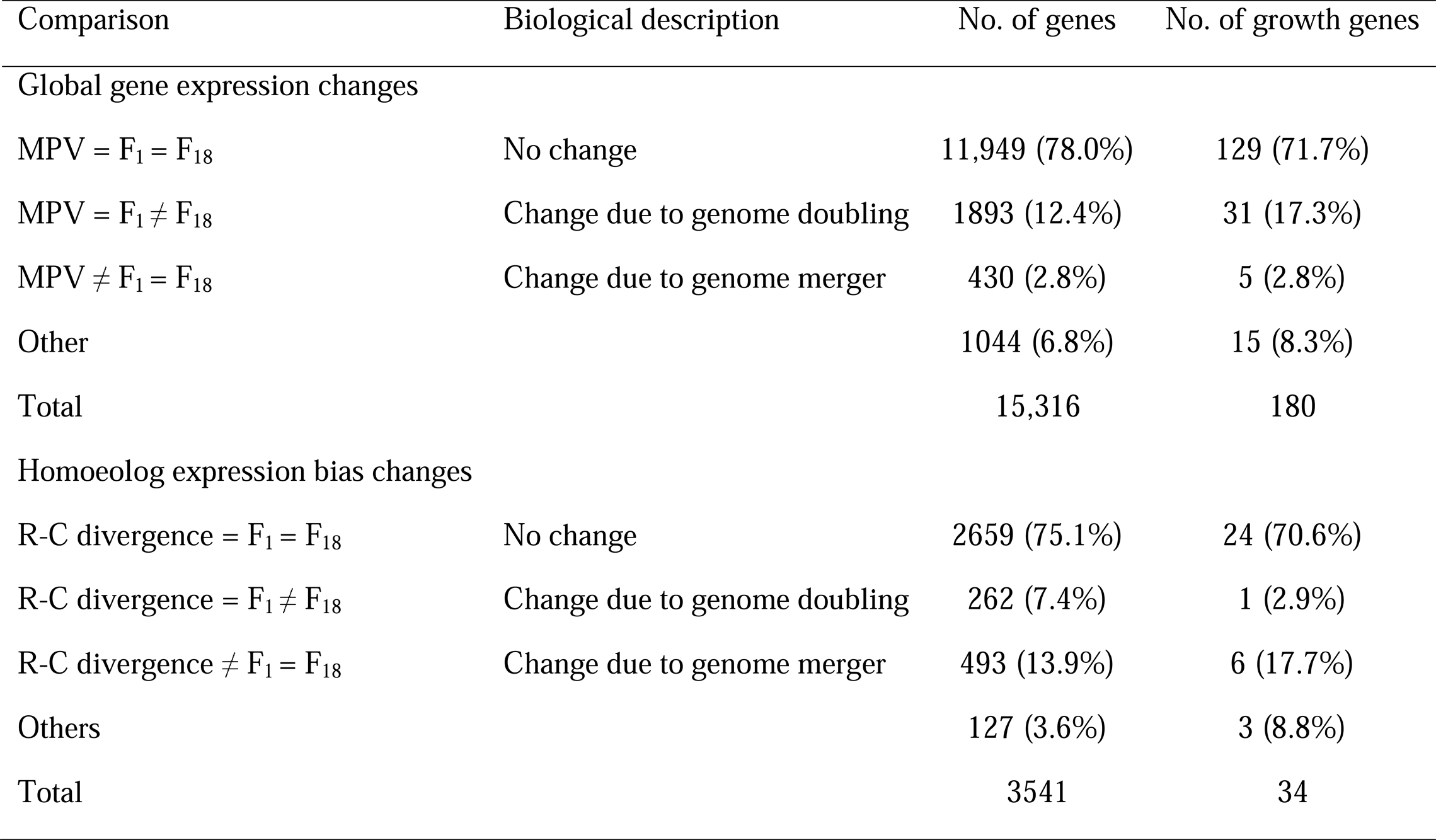
Comparison of gene expression changes and homoeolog expression bias in response to genome merger, genome doubling in F_1_ and F_18_

### The expression pattern of growth-regulated genes using RNA-seq

To investigate how hybridization and polyploidization affect the growth regulatory mechanism in different ploidy level individuals, we used RNA-seq and qPCR to detect HEB in the allotetraploid line of *C. auratus* red var. × *C. carpio.* To analyse the 180 growth-regulated genes, we used the 12 categories of expression patterns to obtain the information on the differential regulation between the hybrids and both parents (up: down = 6: 1 in F_1_, up: down = 2: 8 in F_18_) (*P* = 0.015 in comparisons; Fisher’s exact test) (Figure 2). These results reflected a growth-regulated mRNA preference toward upregulation in the F_1_ and downregulation in the F_18_ compared with the parents. Additionally, we examined per cent of growth-related genes in categories VII and VIII and per cent in categories IX and X. As a result, R-HEB was observed in the F_1_, and C-HEB in the F_18_ (Figure 3).

To further investigate the regulation of HEB related to growth function, all 34 growth-regulated genes were collected from the 3540 genes under HEB analysis (Table 3). Some categories had no statistical significance because of the number of genes selected was a small percentage of the total. However, similar ratios were shown in the other categories. Ultimately, only four R/C-biased growth-regulated genes were identified in the F_1_ and F_18_ (Figure 4). Additionally, a similar situation was observed in the analysis of their expression patterns, in which the value of *silico* MPV was used as a reference point in comparisons with hybrids (Table 4). Among the 180 growth-regulated genes, 71.7% exhibited no expression change in both the F_1_ and F_18_ (Table 4). Thus, global expression and homoeologue expression analysis of growth-regulated genes provided an insight into how changes in expression levels were induced by genome doubling or genome merger and the underlying regulation mechanism.

### Determination of homoeologue expression bias in seven genes using qRT-RCR

To validate whether the patterns of HEB observed above reflected the growth regulation in the F_1_ and F_18_, we detected the HEB of seven key growth-related genes (*igf1, igf2, ghr, tab1, bmp4* and *mstn*) in three tissues (liver, muscle and ovary) using homoeologue-specific qPCR. Interestingly, two scenarios were observed: (1) the silencing of the C homoeologous transcripts of the *mstn* gene was detected in the liver of the F_1_ and F_18_ and the muscle of the F_18_ (Figure 5). (2) Different degrees of HEB were observed in the three tissues (Figure 6). However, R-HEB was observed in most tissues in the F_1_ and F_18_. Compared with the RNA-seq results, homoeologue expression was only verified for the *igf2* genes using qPCR. The results did show similar HEBs between the two methods (Figure 6 and Table S4).

**Figure 5.**
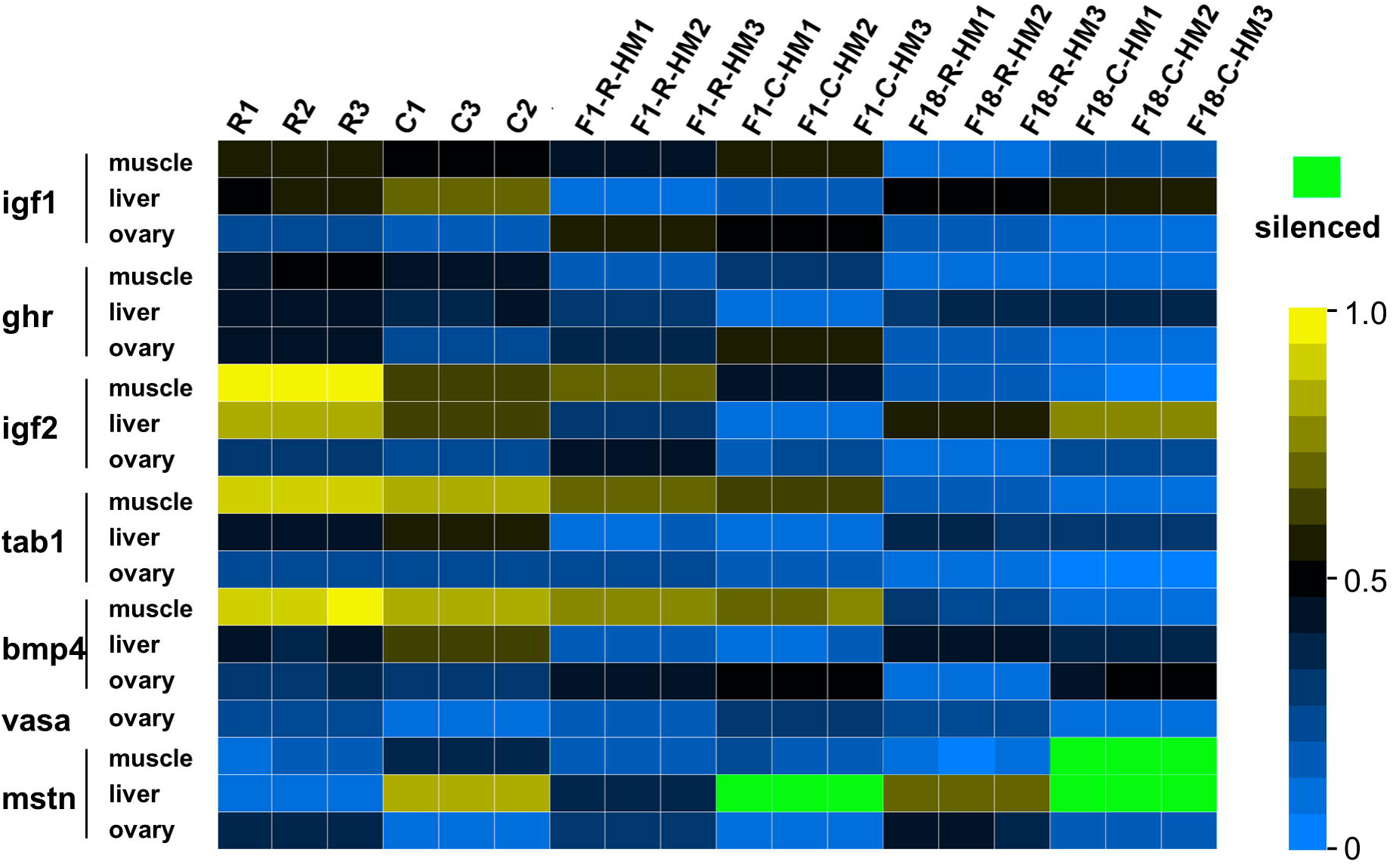

**Figure 6.**
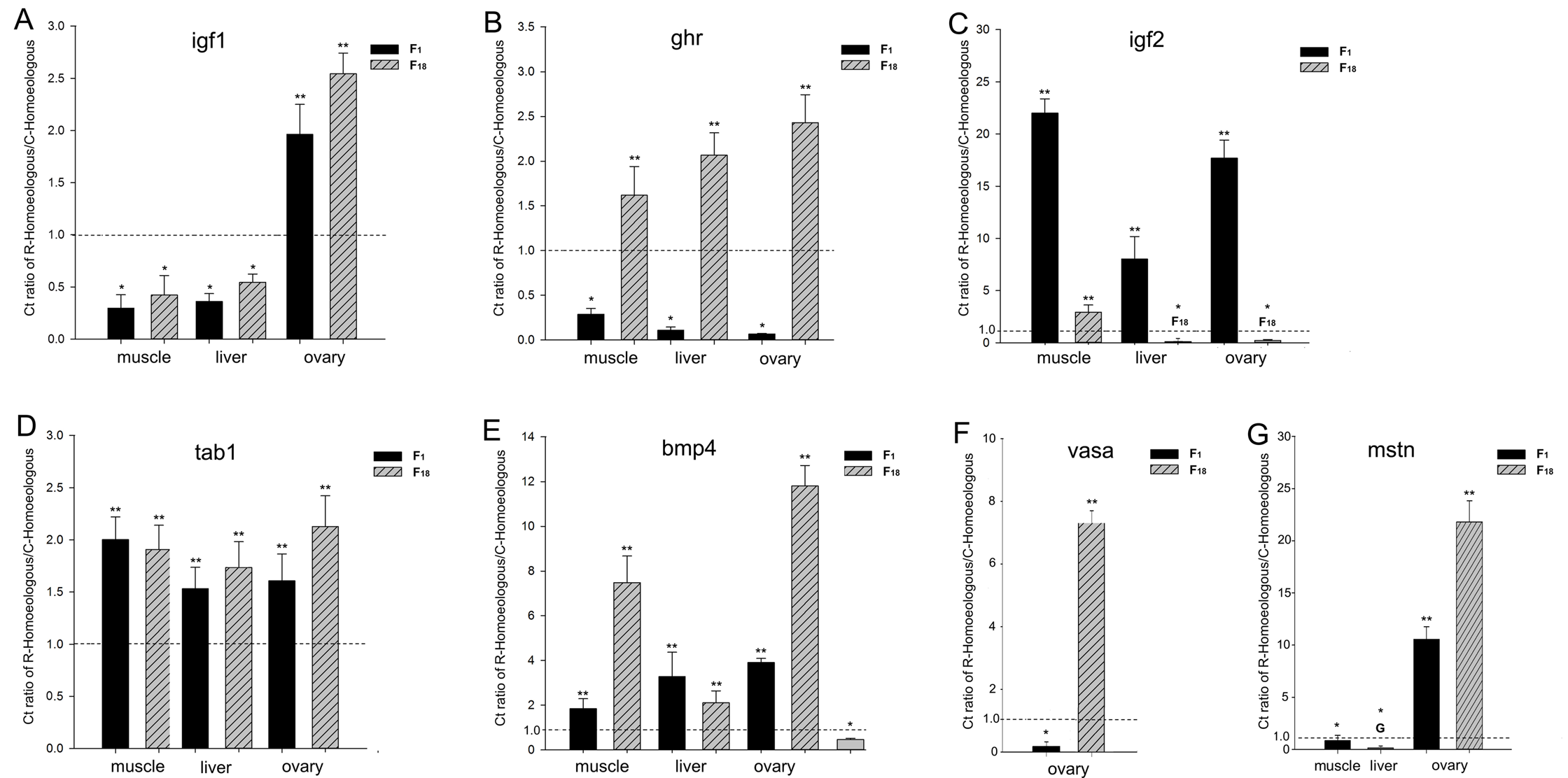

The R to C homoeologue expression level ratio suggested that HEB existed in the different hybrids. We used the ratio to classify the seven homoeologues in the three tissues (Figure 6). For example, C-HEB of the *igf1* gene was detected in the ovary and R-HEB was detected in liver and muscle (Figure 6A). R-HEB of the *ghr* gene was observed in the F_18_, but the F_1_ showed C-HEB (Figure 6B). Interestingly, silencing of C homoeologue expression was observed in the liver of the F_1_, and liver and muscle of the F_18_, which represented overall R-HEB in the progeny. Overall, the phenomenon of R-HEB was obvious in the F_1_ and F_18_ (Figure S3). The expression levels of the R and C homoeologues allowed us to determine how the genetic effect from either of the parents affected F1 and F_18_ (Figure S4).

## Discussion

In this study, distinct genomes of *C. carpio* (C homoeologue) and *C. auratus* red var. (R homoeologue) were merged through hybridization in the F_1_ diploid hybrid, while the F_18_ allotetraploids represented the genome doubling of the F_1_ (Liu *et al*. 2001; Yan *et al*. 2005; Liu *et al*. 2007). Here, we used two approaches (RNA-seq and qPCR) to study the ELD and HEB for total genes and growth-related genes. Our results demonstrated that a decrease in unbalanced ELD and more HEB accompanied hybridization and polyploidization, respectively. The evolution of global expression and R/C homoeologue expression was accompanied by increased HEB and novel expression, as well as increasing levels of silencing of homoeologue expression. A similar analysis was performed on growth-related genes to investigate the relationship between the regulation of growth and homoeologue expression, which provided an insight into growth heterosis under the effect of genome merger and doubling, respectively.

As to the two genomes of the different genera were merged into one cell nucleus, the expression level status from either parent was destroyed. The new expression levels were described as the ELD, where the global expression level resembles that of one of the two parents. Our results demonstrated that the average change in expression level was 22.38% in the F_1_ (*vs*. R = 18.31% and *vs*. C = 26.45%) (Figure 2). After the two types of genome merged, most gene expression levels maintained a steady state. However, the maternal R dominated compared with the paternal D. This phenomenon is frequently observed in hybrid fish, including hybrid *Megalobrama amblycephala* × *Culter alburnus* (Zhou *et al*. 2015), hybrid *Oncorhynchus mykiss* (White *et al*. 2013) and hybrid *Salmo salar* (Debes *et al*. 2012). The new expression levels of the F_1_ were close to MPV (Figure 2). The similar expression levels provided an insight into the character of the hybrid related to heterozygosity, in which two different alleles from different species cooperate in the control of regulatory function.

The study of homoeologue expression level is also an import way to detect the effect of genome merger (Yoo *et al*. 2013; Zhou *et al*. 2015). The co-regulated expression of R and C homoeologues would result in different functions in the hybrids. A previous report on mRNA and microRNA showed that mid-parent expression rarely occurs in genes related to growth and adaptability (Yoo *et al*. 2013; Zhou *et al*. 2015). Thus, the diversified homoeologue expression benefits the combination of advantageous traits in hybrid individuals. Our result for the F_1_ showed no bias of homoeologue expression in 13.9% genes (Table 3), while the majority of genes obtained either of the parental traits after the genome merger. In addition, 15.7% of homoeologue-specific SNPs genes were categorized as overall R/C-biased in F_1_ (Table 3), represent the heterozygosity in most of traits in the hybrid.

The F_18_ allotetraploid is considered as suitable material to study the ELD and HEB under polyploidization, while the genome doubling occurred in F_1_ diploid hybrids. Changes in the expression levels of 3502 (25.5%) genes were identified in the comparison between the F_18_ and the F_1_, which suggested that genome doubling alters the transcriptome more than genome merger. However, comparing the hybrid expression with both of the parents, we detected 18.9% genes as having significant differences in expression in the F_18_ compared with 22.3% in the F_1_ (Figure 2). This suggested that the pattern of expression levels after the genome doubling had been rebuilt. However, the changes in F_18_ did not simply originate from accumulation of genome merger and genome doubling. To address the dimension of expression evolution, we compared *in silico* MPV expression levels to those actually observed in the F_1_ (9.6%) and F_18_ (15.1%). Our analysis showed that the change in global expression in the F18 represented the combined effects of genome doubling and genome merger. Meanwhile, our result showed that the R-ELD in the F_1_ transform to C-ELD in the F_18_ (Figure 3), in contrast to the results for HEB (Table 4). A similar study showed the same trends in polyploid cotton (Yoo *et al*. 2013). These results suggested the reasonable conclusion that genome merger plays the dominant role in the changes in HEB compared with global expression analysis, which was mostly affected by genome doubling. In terms of the scope of transcriptome alterations, we suspect that most changes in gene expression reflect the downstream consequences of the regulatory networks that subtly responded to the stress of the merger of doubling process.

Allopolyploid fish are distributed worldwide and result from artificial or natural selection. Upon crossing the interspecies barrier, the newly formed progeny always display heterosis, such as rapid growth. For the allotetraploid line of *C. auratus* red var. × *C. carpio*, rapid growth was observed in hybrid offspring compared with both parents (Figure S1). However, there has been no study on the underlying mechanism related to growth heterosis. Recent studies have focused on ELD and HEB to analyse the regulation pattern and their underlying mechanisms (Rapp *et al*. 2009; Yoo *et al*. 2013; Zhou *et al*. 2015). These findings show that allelic interactions and gene redundancy result in heterosis in allopolyploids relative to non-coding RNA, DNA, methylation and transcriptome changes (Michalak 2009; Ng *et al*. 2012). In contrast to global expression analysis in teleost hybrids (Liu *et al*. 2012; Zhong *et al*. 2012), the study of homoeologue expression is a promising method to determine the regulation of growth heterosis (Zhong *et al*. 2012).

In the RNA-seq analysis on 118 growth-related genes in the hybrids compared with the MPVs (*in silico*), the study of global expression suggest that 10.0% of growth-related genes in the F_1_ were upregulated, which was higher than that in the F_18_ (3.0% in total genes) (Figure 2). Moreover, the expressions of growth-related gene were downregulated in 10% in the F_1_, which was lower than that in the F_18_ (18.3% of total genes) (Figure 2). In addition, the differential expression analysis between the F_1_ and F_18_ not only suggested that the effects of genome doubling and genome merger cooperate to form a new pattern of growth regulation in the hybrid populations, but also showed that genome doubling resulted in a reduction in growth-regulated gene expression. Previous studies on homoeologous genes support this non-additive expression after genome doubling in allopolyploid wheat (Pumphrey *et al*. 2009) and fish, including carp (Zhou *et al*. 2015), salmon (Pala *et al*. 2008) and cichlid (Albertson and Kocher 2005). The differentially expressed genes between the F_1_ and F_18_ were placed in 12 categories of expression patterns: upregulated (IV, V and VI) and downregulated (I, II, III) growth genes contributed to the lower expression level of homoeologous transcripts in allotetraploids (Figure 3). This result might provide an insight into the rapid growth in the F_1_ compared with the F_18_ (Figure S1).

Maternal-specific expression is observed not only in hybrid plants, but also in lower vertebrates (McKeown *et al*. 2011; Michalak 2014). In the analysis of the categories of growth-related homoeologous genes, the analysis of HEB provided an insight into effect of originating from either of maternal R or paternal C, respectively. The analysis of overall bias identified four genes (*pdgfaa*, *igfbp2a, igfbp1a* and *igfbp1a*) from the 34 homoeologue-specific growth-related genes. The result of R bias analysis in the F_1_ (R *vs*. C = 4.0 *vs*. 0) and F_18_ (R *vs*. C = 3.0 *vs*. 1.0) suggested that homoeologue expression of maternal R plays a major role in the liver transcriptome (Figure 4). Compared with maternal R, the rapid growth characteristics were detected in paternal C. Meanwhile, the joint expression of R/C homoeologues of *igf1* and *ghr* increases the expression diversity and play an import role in promoting the growth ratio in the hybrids (Zhong *et al*. 2012). However, our results for *igf1, igf2* and *ghr* suggested that C-HEB might contribute to rapid growth. Meanwhile, other key growth-related genes (tab1, *bmp4 mstn and vasa*) were used to detect R-/C- HEB (Figure 6), in which regulation of growth was accompanied by different levels of R/C-homoeologue bias. In the R/C bias analysis, although few significant differential homoeologue expression genes were detected in our study, the consequence of potential R-biased was still identified in the analysis of 34 homoeologue-specific growth-related genes (Table 3). The biases of homoeologue-specific genes observed here suggested a role for epigenetic modulation in growth. This phenomenon suggested that the changes in homoeologue expression might contribute to enhance growth and accelerated body development.

Interestingly, silencing of C homoeologues was observed for the growth-related gene *mstn* (Figure 5). One explanation for this observation could be genomic imprinting, implying that gene expression control would be mediated by one parental genome, whereas the genetic material inherited from the other parents is silenced in the hybrid (Martin and McGowan 1995). Some genes always exhibit single-genome-mediated expression in hybrids (McGowan and Martin 1997). A recent study demonstrated that mutations in the *mstn* gene resulted in increased muscle mass and strength in vertebrates, making these individuals considerably stronger than their peers (Schuelke *et al*. 2004). The observation that larger individuals are always seen in hybrid fish populations supports these findings (Shen *et al*. 2006; Yu *et al*. 2011). However, further study is necessary to verify the homoeologue silencing and its relationship with epigenetic traits associated with genome merger and genome doubling.

## Concluding Remarks

Various patterns of global expression and homoeologue expression accompanied by hybridization and polyploidization influence characteristics that are associated with major changes at the epigenetic level, such as growth heterosis. Here, we focused on the total and growth-related transcripts in diploid hybrids and allotetraploids. Our result has indicated that the conversion of homoeologue ELD and the increasing degree of deviation in HEB occurred in the different stages of hybridization and polyploidization. The results for growth-related genes revealed that different degrees of R-HEB contributed to the formation of new growth regulation patterns, and accelerated the genetic diversity of the hybrid population. Global expression and homoeologue expression analysis provided an in-depth understanding of the shape of the expression regulation mechanisms driven by hybridization and polyploidization. In addition, the stability and changeability of growth regulation associated with hybridization and polyploidization provided an insight into the underlying mechanisms.

## Authors’ Contributions

LR carried out bioinformatics analyses and wrote the manuscript. SJL contributed to the conception and design of the study. WHL, XJT and JX provided assistance extracting the raw material. CCT, MT, and CZ modified the manuscript. All authors read and approved the final manuscript.

## Acknowledgements

This work was supported by the Major international cooperation projects of the National Natural Science Foundation of China (Grant No. 31210103918), Key Item of National Natural Science Foundation of China (Grant No. 31430088), Training Program of the Major Research Plan of the National Natural Science Foundation of China (Grant No. 91331105), the National Key Basic Research Program of China (Grant No. 2012CB722305), the Doctoral Fund of Ministry of Education of China (Grant No. 20114306130001), the National High Technology Research and Development Program of China (Grant No.2011 □ AA100403), the Cooperative Innovation Center of Engineering and New Products for Developmental Biology of Hunan Province (20134486) and the construct program of the key discipline in Hunan province and China.

